# Aged Tendons Have Impaired Mechanosensitivity and Lower Thresholds for Injury under Dynamic Compression

**DOI:** 10.64898/2026.04.23.720423

**Authors:** Samuel J. Mlawer, Brianne K. Connizzo

## Abstract

Rotator cuff tendinopathy is highly prevalent in aging populations, yet the mechanisms leading to age-dependent tendon degeneration are not well understood. In addition to tensile loading, tendons are subjected to compressive forces at certain anatomical sites (e.g., Achilles, rotator cuff), where altered adaptive responses may contribute to degenerative remodeling. The objective of this study was to investigate age-related differences in tendon responses to dynamic compressive loading using an *ex vivo* model. Murine flexor tendon explants from young and aged animals were cultured in a biaxial bioreactor and subjected to different levels of dynamic compressive loading. We then observed changes in metabolic activity, matrix composition, matrix biosynthesis, matrix structure, and gene expression. Young tendons exposed to moderate levels of compression maintained homeostasis, whereas high compression induced a robust adaptive response characterized by increased glycosaminoglycan accumulation, elevated collagen content, and upregulation of remodeling-associated genes including collagen I, decorin, and MMP-9, as well as inflammatory and apoptotic markers. In contrast, aged tendons demonstrated a qualitatively different response, with transcriptional downregulation of key remodeling markers alongside elevated secretion of matrix-degrading enzymes and pro-inflammatory cytokines, indicative of a maladaptive mechanobiological response even at low compressive levels. These findings reveal that impaired mechanosensitivity and a lower threshold for injury may predispose chronically loaded tissues to degenerative pathology associated with excessive compressive loading.

## Introduction

Rotator cuff pathology is increasingly prevalent in aging populations, with rates reaching as high as 80% in individuals over 80 years old (Milgrom et al., 1995). Despite this high burden, the mechanisms that drive tendon degeneration, and how these processes are altered with aging, are not fully understood. Mechanical loading is a key regulator of tendon homeostasis, and while tendons are primarily adapted to resist tensile forces, several anatomical sites are also subjected to compressive loads from adjacent bony structures. The supraspinatus tendon, which passes beneath the acromial–clavicular arch, is a common example of a tendon exposed to repeated compression (Seitz et al., 2011). In regions of regular compression, tendons develop a fibrocartilaginous phenotype characterized by rounded cells and increased expression of cartilage-associated extracellular matrix (ECM) molecules. This phenotype helps buffer compressive stresses by increasing proteoglycan-mediated hydration. However, when areas that are predominantly designed for tensile loading, or transition zones between fibrocartilaginous and midsubstance regions, are repeatedly exposed to abnormal or excessive compression, their inability to accommodate these forces can initiate degenerative changes and accelerate pathology (Benjamin & Ralphs, 1998). Age-related alterations in matrix composition, cellularity, and mechanosensitivity may further impair the ability of tendon to adapt to compressive loading, potentially accelerating degenerative remodeling under chronic mechanical stress (Cuomo et al., 1998; Kwan et al., 2023).

External compression has therefore been identified as an important contributor to tendinopathy, yet relatively few studies have examined the direct biological responses of tendon to controlled compressive loads or how these responses differ with aging (Carpenter et al., 1998; Seitz et al., 2011). To address this gap, we leveraged our tendon explant model that preserves native ECM organization and resident cells with the design of a novel multiaxial loading bioreactor to facilitate precise application of tensile and compressive loading simultaneously on whole tissues (Aggouras et al., 2024). Using this platform, our prior work demonstrated that a single acute compressive overload induced proteoglycan turnover and inflammatory signaling in young tendons, whereas aged tendons exhibited minimal transcriptional or matrix responses, suggesting impaired mechanosensitivity (Mlawer et al., 2023). Because acute loading alone was insufficient to stimulate a robust response in aged tissue, we next sought to determine whether continuous dynamic compression could drive age-dependent adaptive or degenerative responses.

Therefore, the purpose of this study is to explore age-related differences in the response of tendon explants to continuous dynamic compressive loads. We hypothesized that young tendons would exhibit a robust remodeling response characterized by increased ECM synthesis, whereas aged tendons would demonstrate a blunted or dysregulated response consistent with impaired mechanosensitivity. We further hypothesized that higher compressive loading would promote a shift toward a fibrocartilaginous, injury-associated phenotype accompanied by decreased collagen fiber alignment, inflammation, and degeneration.

## Methods

### Explant Harvest and Culture

Directly following sacrifice, flexor digitorum longus (FDL) tendon explants were harvested from the hind limbs of young (4 months; n=32 animals) and aged (22-24 months; n=36 animals) C57BL/6J male mice using previously described methods per approved animal use protocol (BU IACUC PROTO202000046). After harvest, explants were loaded into grips with a 10-mm gauge length and placed into a custom-built multiaxial loading bioreactor (Figure 1A-B).

**Figure 1.**
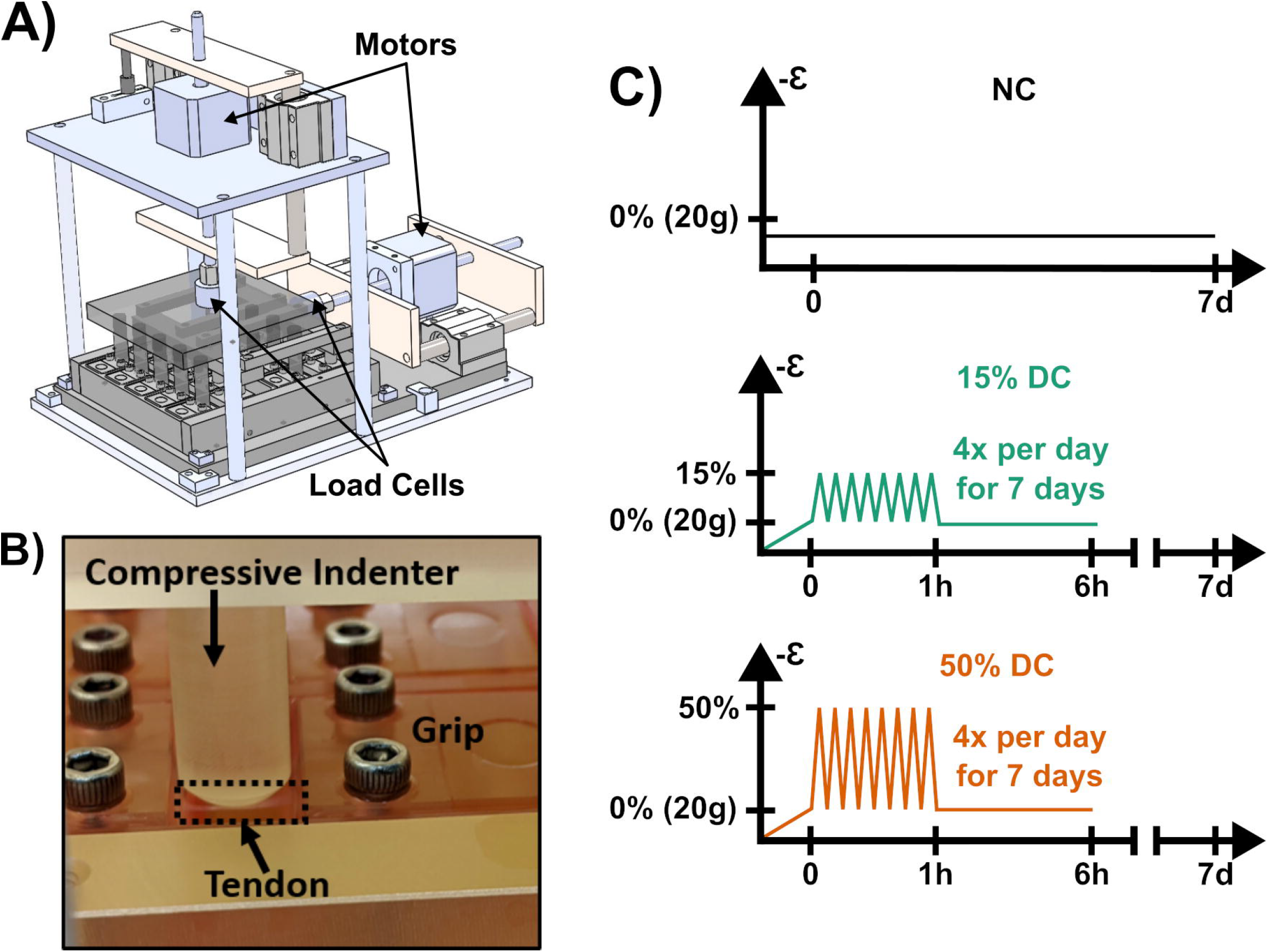
(A) Biaxial loading device used to maintain tension while inducing acute compressive injury at D0 with (B) a close-up of a single well showing a tendon being compressively loaded. (C) Three loading protocols used in this study: non-compression (NC), 15% dynamic compression (15% DC), or 50% dynamic compression (50% DC). All groups maintained under 3% static tension.

Throughout culture, explants were kept in culture medium consisting of low glucose Dulbecco’s Modified Eagle’s Media (1 g/l; Fisher Scientific) supplemented with 10% fetal bovine serum (Cytiva, Marlborough, MA), 100 units/mL penicillin G, 100 µg/mL streptomycin (Fisher Scientific), and 0.25 µg/mL Amphotericin B (Sigma-Aldrich).

### *Ex Vivo* Mechanical Loading

After being placed in the bioreactor, all explants were loaded to 3% static tensile strain at 1% strain/sec and held at this tension for the duration of the experiment. Following tensioning, explants randomly assigned to the 15% and 50% dynamic compression groups were subjected to a 7-day dynamic compression protocol, consisting of a 1-hour displacement-controlled cyclic loading (15% and 50% strain respectively, 1 Hz) followed by 5-hours without compression. This 6-hour cycle was repeated four times per day throughout the culture period. The non-compression (NC) group experienced only static tensile strain for 7 days.

### Metabolism, Biosynthesis, and Composition

Explant metabolic activity was measured using a resazurin reduction assay, as previously described (Rodríguez-Corrales & Josan, 2017). Briefly, explants were incubated for 3 hours in culture media containing resazurin solution, after which the fluorescence intensity of the reduced metabolite, resorufin, was measured on a microplate reader. Measurements were normalized to same-day control wells containing only media. Following culture, explants (n = 9-14/group) were rinsed for 2 hours and then digested overnight (16 hours) in proteinase K (5 mg/mL; Sigma-Aldrich). Sulfated glycosaminoglycans (sGAG) and total protein synthesis (as an indicator of collagen synthesis) were quantified by 24-hour incorporation of ^35^S-sulfate (20 μCi/mL) and ^3^H-proline (5 μCi/mL), respectively (Perkin□Elmer). Radioactivity in the digested tissue was measured with liquid scintillation counter (Perkin-Elmer) (Connizzo et al., 2020).

Double□stranded DNA content was quantified using the PicoGreen fluorescence assay (Ahn et al., 1996). sGAG content in the tissue digest was determined using the dimethylmethylene blue (DMMB) assay (Farndale et al., 1986). For collagen content, 100μL of each digest was hydrolyzed in 12 M HCl, dried, reconstituted, and analyzed using the hydroxyproline (OHP) assay (Dourte et al., 2013). All biochemical data were normalized to tendon dry weight.

### Histology

Following culture, explants (n=4/group) were embedded in optimal cutting temperature (OCT) compound (Fisher Scientific) and 10 µm thick cross sections were collected on glass slides. Prior to staining, sections were fixed using 4% paraformaldehyde (Electron Microscopy Services). For hematoxylin and eosin (H&E) staining, slides were immersed in hematoxylin (Harris Hematoxylin; Electron Microscopy Sciences) for 4 min and eosin (Electron Microscopy Sciences) for 10 seconds. For toluidine blue staining, slides were immersed in toluidine blue (Fisher Scientific) for 1 min while being gently agitated up and down. Both H&E and toluidine blue slides were imaged at 20x with a VS120 virtual slide scanner (Olympus Scientific Solutions, Waltham, MA). H&E images were evaluated using a custom MATLAB script for cell density and cell aspect ratio. Toluidine blue images were graded using a custom MATLAB script quantifying the proportion of purple hue. Second harmonic generation (SHG) images were obtained with a Bruker Investigator two-photon microscope with 1040 nm excitation laser and 525 ± 70 nm bandpass detection filter for SHG detection. SHG images were evaluated for collagen z alignment using FiberFit software from Boise State University,(Morrill et al., 2016).

### Gene Expression

Explants collected at Day 0 and Day 7 (n = 5–7 per group per day) were flash-frozen in liquid nitrogen and stored at −80 °C until RNA isolation. Tissues were homogenized in TRIzol reagent (Invitrogen) using a bead homogenizer (Benchmark Scientific). The resulting aqueous phase was purified using the Quick-RNA purification kit (Zymo Research) following the manufacturer’s protocol. Complementary DNA (cDNA) was synthesized from the isolated RNA via reverse transcription, and quantitative PCR (qPCR) was performed using a StepOnePlus Real-Time PCR System (Applied Biosystems, Foster City, CA). Gene expression levels were calculated using the ΔΔCT method, normalized to the housekeeping gene β-actin and to Day 0 control values.

### Secreted Protein Analysis

Protein levels of matrix metalloproteinases (MMPs; MMP-3, MMP-9, MMP-13) were determined using a custom multiplex ELISA. MMP-9 was detected using the U-PLEX Mouse MMP-9 (total) antibody set and MMP-3 was detected using the R-PLEX Mouse MMP-3 (total) antibody set (Meso Scale Discovery). The MMP-13 antibody set was purchased separately (Abcam; ab314458), conjugated in-house, and validated using a recombinant protein calibrator (Cusabio; CSB-EP014660MO; 20,000 pg/mL). The custom multiplex assay was performed per manufacture’s guidance using the U-PLEX standard protocol. Protein levels of inflammatory cytokines (IL-1β, IL-4, IL-6, IL-10, IL-15, TNFα, KC/GRO, MCP-1) were determined using a U-PLEX Custom Biomarker Assay kit (Meso Scale Discovery). Protein analyses were determined via analysis of spent culture medium (n=4-6/group).

### Statistics

Data points exceeding two standard deviations from the group mean were classified as outliers and excluded from analysis. Statistical analyses were conducted using GraphPad Prism 8 (GraphPad Software, San Diego, CA). Differences among loading groups were assessed by one-way analysis of variance (ANOVA) followed by Bonferroni corrected t-tests. Statistical significance was defined as p < 0.05, and all results are reported as mean ± 95% confidence interval.

## Results

We first assessed how dynamic compression influenced markers of the most common ECM component, collagen. Collagen content increased only in young tendons subjected to 50% DC (Figure 2A). This increase occurred without corresponding changes in collagen synthesis.

**Figure 2.**
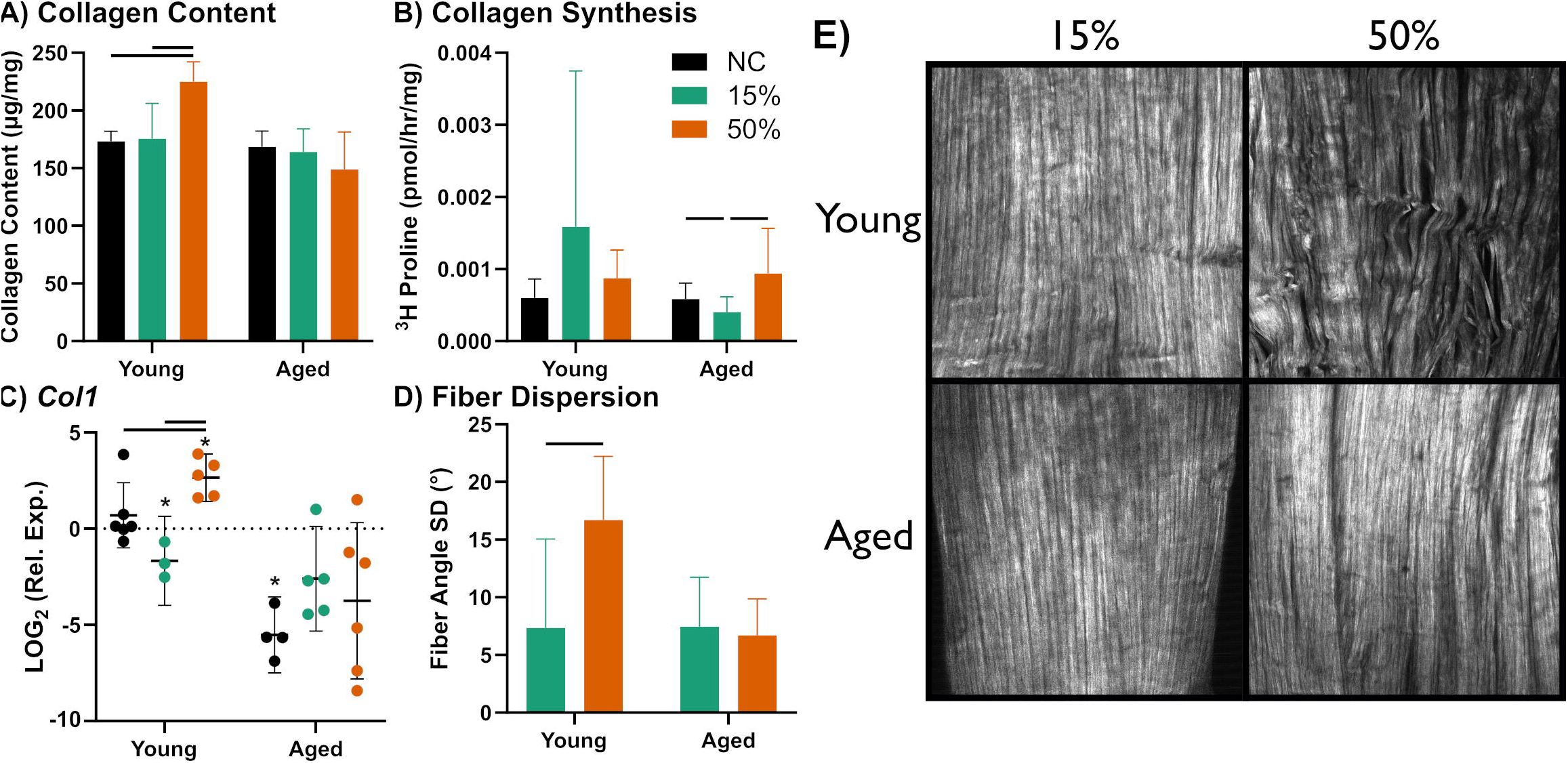
Measurements of (A) collagen content, (B) collagen synthesis, (C) collagen I gene expression, (D) fiber dispersion quantified from second harmonic generation (SHG) imaging. (E) Representative SHG images for each group. Data is presented as mean ± 95% confidence interval. Significance between groups is denoted with a bar (-) with p<0.05.

Aged tendons exposed to 15% DC exhibited reduced collagen synthesis compared to both NC and 50% DC (Figure 2B). In young tendons, expression of collagen I was upregulated by 50% DC (Figure 2C). There were no differences with compression in aged groups, although it is noted that collagen expression was markedly lower in aged compared to young tendons in all loading groups. Finally, we used SHG to examine the collagen structure and found that 50% dynamic compression decreased collagen fiber alignment in young tendons (Figure 2D,E).

We next evaluated how another key ECM component, proteoglycans and their associated sulfated glycosaminoglycan (sGAG) chains, were altered in response to compressive loading. In aged explants, 50% DC increased sGAG content relative to 15% DC (Figure 3A). Although sGAG content remained unchanged in young tendons, sGAG synthesis was elevated at 50% DC (Figure 3B). For proteoglycan-related genes, decorin expression decreased with 15% DC but increased with 50% DC in young explants (Figure 3C). Biglycan expression declined at both loading levels in young tendons and at 15% DC in aged tendons (Figure 3D). Fibromodulin expression decreased with 15% DC in young samples but increased with 50% DC in aged explants (Figure 3E). Expression of the cartilage-associated proteoglycan, aggrecan, increased from 15% to 50% DC in aged samples only (Figure 3F). We also used toluidine blue to stain for GAGs and found no differences in metachromatic staining. However, an increase in blue nuclear staining was observed (Figure 3G–H).

**Figure 3.**
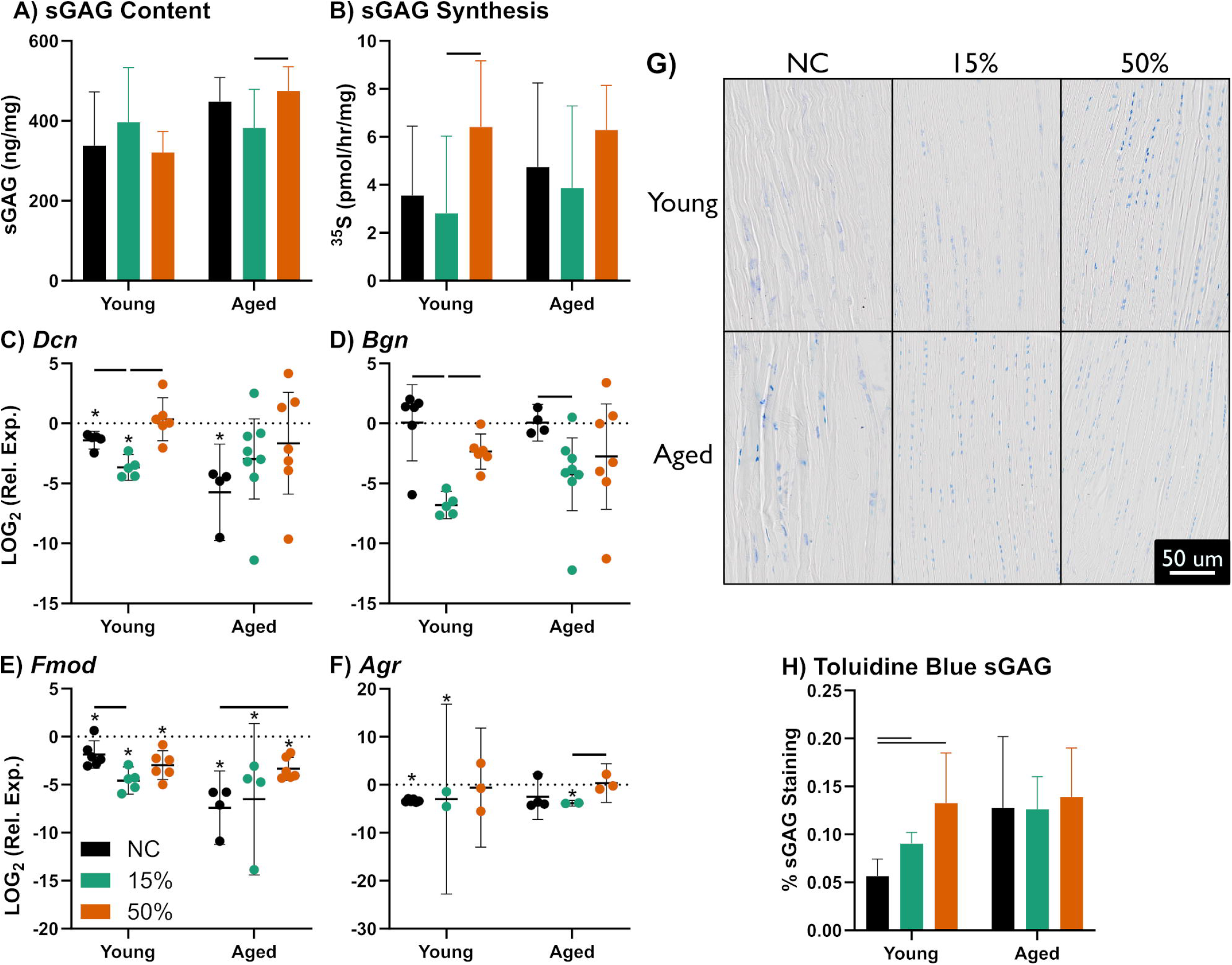
Measurements of (A) sGAG content, (B) sGAG synthesis, as well as gene expression of (C) decorin, (D) biglycan, (E) fibromodulin, (F) and aggrecan. (G) Representative toluidine blue images and (H) quantification via custom image-based processing. Data is presented as mean ± 95% confidence interval. Significance between groups is denoted with a bar (-) with p<0.05.

We then examined markers of collagen and proteoglycan degradation. While MMP-3 expression was unaffected by loading in all groups, we found an increase in MMP-3 protein secretion in the media from aged tendons between NC and 15% DC (Figure 4A). We found that MMP-9 expression increased with 50% DC in young explants but was downregulated in aged tendons; however, these differences were not observed in protein secretion (Figure 4B). In contrast, MMP-13 expression was downregulated by 50% DC in both age groups. For protein secretion, we saw a decrease in MMP-13 at 15% DC in young tendons (Figure 4C). Notably, MMP-9 and MMP-13 were below the limit of detection in non-compressed samples, whereas both compression conditions yielded detectable levels, suggesting loading-induced secretion that is not captured by statistical comparisons (Figure 4B,C).

**Figure 4.**
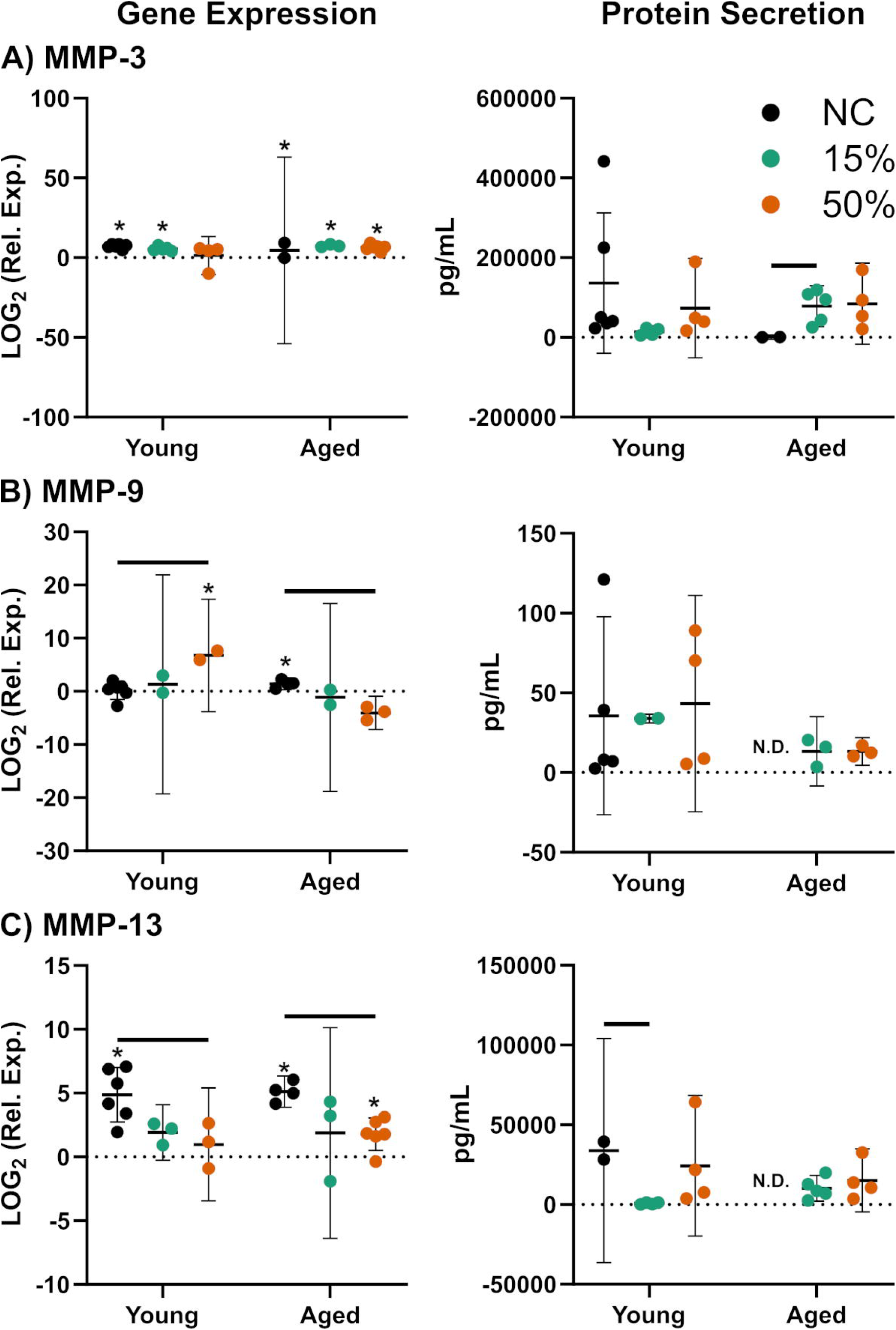
Measurements of gene expression (left) and protein secretion (right) of (A) MMP-3, (B) MMP-9, (C) and MMP-13. Data is presented as mean ± 95% confidence interval. Significance between groups is denoted with a bar (-) with p<0.05. N.D. denotes non-detected groups.

Next, we used H&E staining to see how these ECM changes affected cell morphology. We found that in both age groups, compressive loading decreased cell density (Figure 5A,B). Nuclear aspect ratio increased with loading in young tendons only, indicating a rounding of cells with compression (Figure 5A,C). We then assessed how compressive loading affected markers of cellular metabolism and injury. In young explants, 50% DC reduced metabolic activity (Figure 6A). DNA content, however, was decreased only in aged tendons subjected to 15% DC (Figure 6B). For gene markers of injury, IL-6 was upregulated by compression in young tendons (Figure 6C). IL1β was downregulated at both loading levels in aged tendons (Figure 6D). TNF-α decreased only with 15% DC in young tendons (Figure 6E). Finally, we found that caspase-3 was upregulated by 50% DC in young tendons but unchanged in aged tendons (Figure 6F).

**Figure 5.**
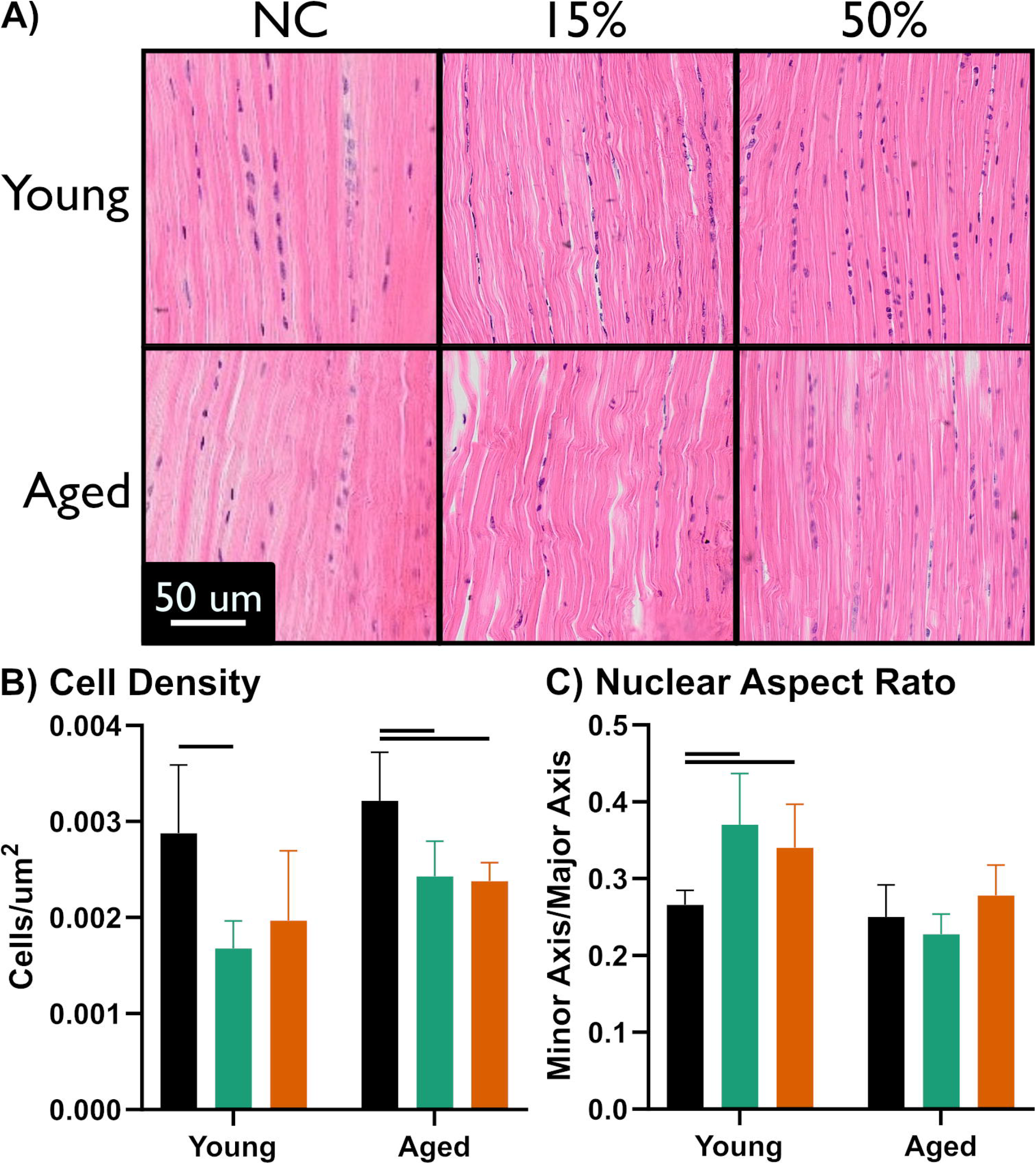
(A) Representative images of H&E staining, with quantifications of (B) cell density and (C) nuclear aspect ratio from the images. Data is presented as mean ± 95% confidence interval. Significance between groups is denoted with a bar (-) with p<0.05.

**Figure 6.**
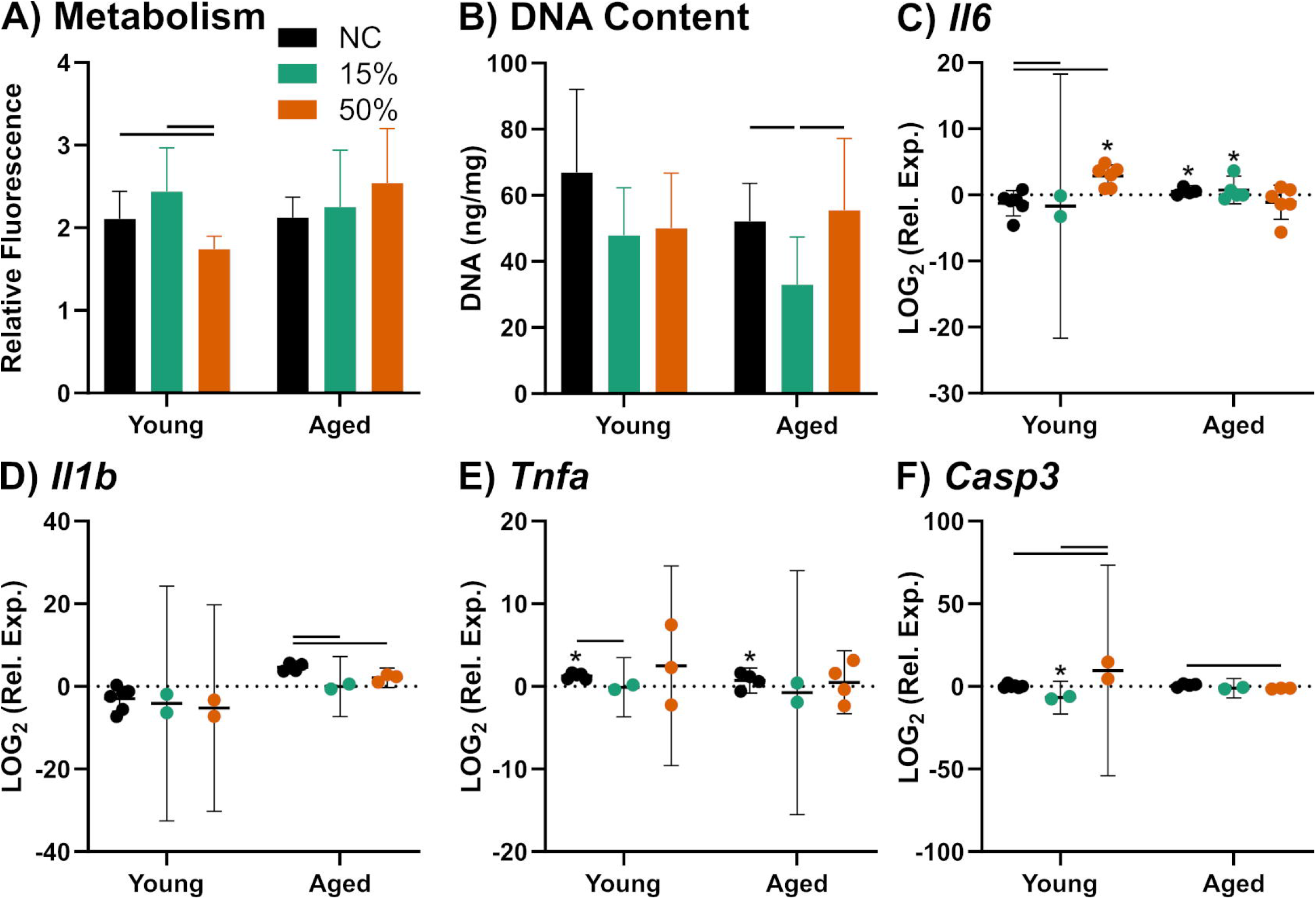
Measurements of (A) metabolic activity, (B) DNA content, and gene expression of injury and apoptosis markers (C) IL-6, (D) IL-1β, (E) TNF-α, (F) and caspase-3. Data is presented as mean ± 95% confidence interval. Significance between groups is denoted with a bar (-) with p<0.05.

When we looked at protein levels of inflammation markers in young tendons we observed decreases in IL-4, IL-15, TNF-α, KC/GRO, and MCP-1 with 15% DC. However, except for KC/GRO, these differences disappeared when loading increased to 50% DC (Figure 7A,C,F,G,H). Interestingly, this pattern was not found in IL-10, IL-6, or IL-1β (Figure 7B,D,E). In aged tendons we found an increase in IL-4 with 15% DC (Figure 7A) and an increase in KC/GRO with both levels of compression (Figure 7G). All other protein levels were not quantitatively different between loading groups in aged tendons. However, for both IL-6 and MCP-1, the aged non-compressed samples were below the limit of detection, while the compressed groups reported detectable values, implying an increase in these proteins with compression that is not quantifiable with current detection limits (Figure 7D,H).

**Figure 7.**
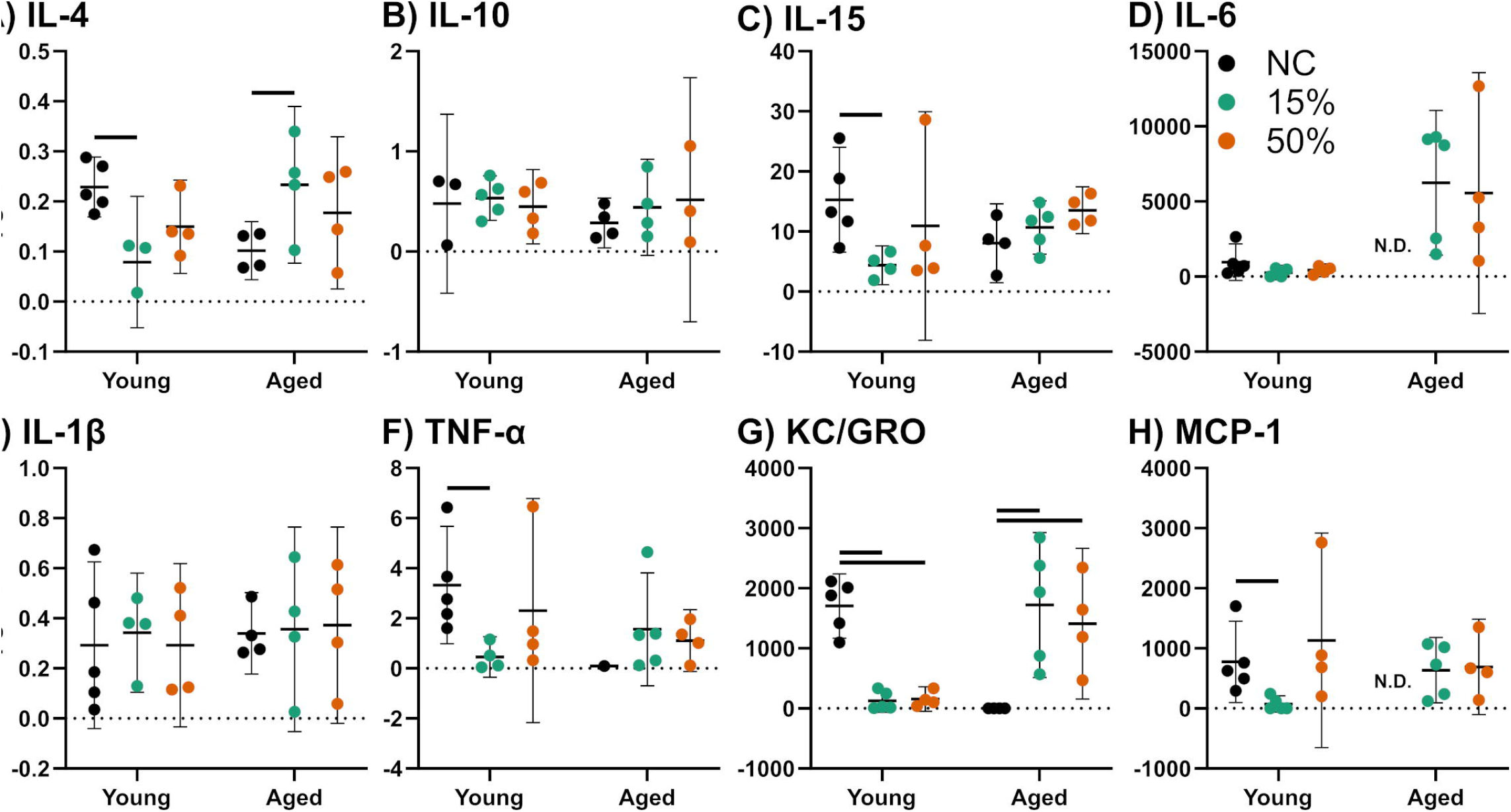
Measurements of protein secretion in media of (A) IL-4, (B) IL-10, (C) IL-15, (D) IL-6, (E) IL-1β, (F) TNF-α, (G) KC/GRO, (H) and MCP-. Data is presented as mean ± 95% confidence interval. Significance between groups is denoted with a bar (-) with p<0.05. N.D. denotes non-detected groups.

## Discussion

In this study, we found that tendon’s responses to dynamic compression are both age- and magnitude-dependent. Similar to previous studies from our lab, our results suggest that young tendons show a much more robust response than aged tendons, which struggle to adapt to their environment (Aggouras et al., 2024; Mlawer et al., 2023, 2025). When young tendons experience the supraphysiological load of 50% dynamic compression, they induce adaptation by increasing sGAG synthesis and expression of collagen. Interestingly, they also upregulate decorin, a regulator of collagen assembly, which results in incorporation of new collagen into the structure, increasing collagen content overall. Expression of MMP-9 was also increased, suggesting collagen-related degradation, perhaps due to damage clearance or structural re-organization.

Together, these changes suggest active extracellular matrix remodeling in response to high compressive loads. Importantly, we also see increases in apoptotic and inflammatory markers, which likely reflect a regulated stress response that facilitates tissue turnover and repair rather than pathological injury, consistent with a conventional tendon repair process (Gracey et al., 2020).

Interestingly, young tendons exposed to the more moderate 15% dynamic compression did not exhibit the same remodeling response observed at 50% strain. Collagen content, collagen I expression, and sGAG synthesis remained unchanged, indicating that this loading magnitude did not stimulate ECM accumulation. Instead, moderate compression was associated with reduced expression of several small leucine-rich proteoglycans along with decreased MMP-13 levels.

Additionally, both pro- and anti-inflammatory markers were downregulated under 15% compression. These findings suggest that moderate compressive loading functions primarily as a homeostatic mechanical stimulus that maintains tissue quiescence while suppressing ECM turnover and inflammatory signaling, rather than triggering active remodeling. This may reflect a threshold-dependent mechanobiological response, where 15% dynamic compression does not exceed the level required to activate remodeling pathways. Similar threshold-dependent responses to mechanical loading have been described in other musculoskeletal tissues, such as bone, where a minimum strain magnitude is required to initiate bone formation (Robling & Turner, 2009). This ability to tolerate physiological levels of compression without initiating injury or repair pathways highlights the capacity of young tendon to discriminate between moderate and supraphysiological mechanical stimuli.

In contrast, aged tendons exhibit a markedly different response to compressive loading, with minimal evidence of adaptive ECM remodeling at either level. When exposed to the same supraphysiological mechanical stimulus, aged tendons showed a downregulation of key remodeling and stress-response markers, including MMP-9, MMP-13, IL-1β, and caspase-3, despite there also being an increase in collagen synthesis and aggrecan expression. Interestingly, analysis of media at day 6 revealed a concurrent elevation of secreted MMP-3, MMP-9, and MMP-13, alongside pro-inflammatory cytokines IL-6, TNF-α, KC/GRO, and MCP-1. This dissociation between transcriptional downregulation of remodeling genes at day 7 and elevated protein secretion of matrix-degrading enzymes and inflammatory mediators at day 6 may reflect the temporal dynamics of the injury response, where gene expression represents a later shift following the initial upregulation shown by protein secretion. Alternatively, it may also suggest a more fundamental uncoupling of ECM synthesis from degradation and injury signaling at the post-transcriptional level. In either case, this pattern could be indicative of a dysregulated mechanobiological response rather than a simple absence of change. Age-associated alterations in mechanosensitivity have been documented across connective tissues, with aged cells and matrices exhibited altered focal adhesions and diminished ability to transduce mechanical stimuli into adaptive biochemical responses (Han et al., 2025; Phillip et al., 2015). Interestingly, we do see a response here, but it is consistent with maladaptive remodeling rather than growth and adaptation. This may predispose chronically loaded aged tendons to matrix disorganization and degenerative changes over time (Gögele et al., 2025; Korcari et al., 2023). Notably, aged tendons also exhibited dysregulated responses under the more moderate 15% loading condition. Rather than maintaining the homeostatic state observed in young tendons, moderate compression in aged samples was associated with reduced collagen synthesis, decreased cellularity, and altered inflammatory signaling. Together, these findings suggest that aging diminishes the ability of tendon to appropriately distinguish between physiological and supraphysiological mechanical stimuli, further supporting the presence of impaired mechanosensitivity.

Histological analyses further supported the age- and load-dependent remodeling observed at the molecular level. SHG imaging revealed decreased collagen alignment in young tendons exposed to 50% dynamic compression. In contrast, collagen alignment was preserved in young tendons subjected to 15% dynamic compression, indicating that moderate loading did not disrupt fibrillar organization. Aged tendons likewise showed no detectable changes in collagen organization under either loading condition, consistent with their limited biochemical response. Previous studies have used SHG to detect reduced collagen alignment and fiber organization in injured or overused tendons (Abraham et al., 2011; Durgam et al., 2020). Despite increased sGAG synthesis in young tendons, we didn’t observe any robust metachromatic staining indicative of a high concentration of GAGs, but rather saw increased blue staining localized near cell nuclei. This pattern suggests active proteoglycan production and early-stage matrix remodeling rather than stable GAG accumulation (Vidal & Mello, 2019). Additionally, increased cell rounding was observed in both young compression groups but not in aged tendons. Since it is not accompanied by an upregulation of Sox9, collagen II, or metachromatic GAG staining, this likely does not indicate a shift towards a fibrocartilaginous phenotype. Instead, this cell rounding likely reflects mechanoadaptation and indicates that tenocyte morphology is sensitive to its loading environment. The absence of cell rounding in aged tendons further suggests that age-related impairments limit the ability of cells to dynamically adjust their morphology in response to mechanical load.

We also compared this study’s results back to our previous work where we looked at the effect of a single compressive overload at 50% strain, as well as to 1-hour loading of 50% dynamic compression on day 0 only (Figure S1) (Mlawer et al., 2023). We found an interesting phenomenon in young tendons where sGAG synthesis was upregulated in all three 50% loading groups, but this only resulted in sGAG content in the 2 shorter term loading groups, indicating that tendon needs adequate recovery time for matrix remodeling, similarly to previous results (Muljadi & Andarawis-Puri, 2023). These findings suggest that the interval between loading bouts is a critical determinant of whether active synthesis translates into net proteoglycan accumulation. When we look at collagen, we find that collagen content only increases significantly in our 50% dynamic compression group, implying that repeated loading is required for collagen accumulation. This is consistent with previous work, which observes turnover in collagen to be much slower than that of proteoglycans (Choi et al., 2020). Together, these results show that different loading and rehab programs must be used in order to promote proper remodeling of different ECM components in tendon.

Several limitations should be considered when interpreting these findings. First, this study used an ex vivo tendon model, which allows for precise control of loading magnitude and duration, but does not fully represent the complex *in vivo* environment, including vascularization, immune cell contributions, and systemic signaling. Second, analyses were conducted at a single post-loading time point, which limits our ability to resolve the full remodeling progression, particularly given that transcriptional changes inherently precede ECM-level responses. Future studies incorporating multiple time points would help clarify the temporal dynamics of ECM remodeling following mechanical stimulation. Third, while the rotator cuff represents a more physiologically relevant site of compressive tendon loading in humans, the small size of the murine rotator cuff makes it incompatible with our loading bioreactor system. The FDL tendon was therefore selected as a well-established model for studying loading mechanics in mice, however future work in larger animal models may better reflect the clinical context of rotator cuff pathology. Finally, this work only examines changes in male mice. Ongoing studies are currently under way looking at the role of sex in the response to compressive loading.

In summary, this study demonstrates that tendon responses to dynamic compression are strongly dependent on both age and loading magnitude. Young tendons exhibited robust, strain-dependent mechanoadaptation characterized by changes in ECM synthesis, content, gene expression, and tissue structure, consistent with active matrix remodeling under compressive load. In contrast, aged tendons showed a muted and dysregulated response, with limited structural adaptation and altered regulation of remodeling and stress-response pathways despite exposure to the same mechanical stimuli. Notably, aged tendons also failed to exhibit the homeostatic tolerance to moderate loading observed in young tendons, suggesting that aging impairs the ability of tendon to appropriately distinguish between physiological and supraphysiological mechanical stimuli. Importantly, our findings suggest that increases in ECM synthesis do not necessarily translate into immediate matrix accumulation, highlighting the requirement for time-dependent remodeling and repeated loading for stable ECM incorporation. Together, our data underscore age-related impairments in tendon mechanosensitivity and adaptive capacity and provide new insight into how compressive loading shapes tendon remodeling. These results may provide insights into the development of age-appropriate mechanical loading strategies for tendon repair and rehabilitation.

## Acknowledgements

This study was supported by Boston University and the National Institute of Health (R00□AG063896, R35-GM151127). Research reported in this publication was supported by the Boston University Micro and Nano Imaging Facility (NIH/NIA S10-OD024993), as well as the Neurophotonics Center at Boston University.

## Author Contributions

**Samuel J. Mlawer:** Conceptualization, Methodology, Software, Formal Analysis, Investigation, Validation, Data Curation, Writing - Original Draft, Writing - Review & Editing, Visualization. **Brianne K. Connizzo:** Conceptualization, Methodology, Resources, Writing – Review & Editing, Supervision, Funding Acquisition.

## Figure Captions

**Figure S-1.** Measurements of (A) sGAG content, (B) sGAG synthesis, (C) collagen content, and (D) collagen synthesis across all loading groups. Data is presented as mean ± 95% confidence interval. Significance between groups is denoted with a bar (-) with p<0.05.

## Notes

### Competing Interest Statement

The authors have declared no competing interest.

## References

Abraham, T., Fong, G., & Scott, A. (2011). Second harmonic generation analysis of early Achilles tendinosis in response to in vivo mechanical loading. BMC Musculoskeletal Disorders, 12, 26. 10.1186/1471-2474-12-26

Aggouras, A. N., Stowe, E. J., Mlawer, S. J., & Connizzo, B. K. (2024). Aged Tendons Exhibit Altered Mechanisms of Strain-Dependent Extracellular Matrix Remodeling. Journal of Biomechanical Engineering, 146(071009). 10.1115/1.4065270

Ahn, S. J., Costa, J., & Emanuel, J. R. (1996). PicoGreen quantitation of DNA: Effective evaluation of samples pre-or post-PCR. Nucleic Acids Research, 24(13), 2623–2625.

Benage, L. G., Sweeney, J. D., Giers, M. B., & Balasubramanian, R. (2022). Dynamic Load Model Systems of Tendon Inflammation and Mechanobiology. Frontiers in Bioengineering and Biotechnology, 10. 10.3389/fbioe.2022.896336

Benjamin, M., & Ralphs, J. R. (1998). Fibrocartilage in tendons and ligaments—An adaptation to compressive load. Journal of Anatomy, 193(Pt 4), 481–494. 10.1046/j.1469-7580.1998.19340481.x

Carpenter, J. E., Flanagan, C. L., Thomopoulos, S., Yian, E. H., & Soslowsky, L. J. (1998). The Effects of Overuse Combined With Intrinsic or Extrinsic Alterations in an Animal Model of Rotator Cuff Tendinosis. The American Journal of Sports Medicine, 26(6), 801–807. 10.1177/03635465980260061101

Choi, H., Simpson, D., Wang, D., Prescott, M., Pitsillides, A. A., Dudhia, J., Clegg, P. D., Ping, P., & Thorpe, C. T. (2020). Heterogeneity of proteome dynamics between connective tissue phases of adult tendon. eLife, 9, e55262. 10.7554/eLife.55262

Connizzo, B. K., Piet, J. M., Shefelbine, S. J., & Grodzinsky, A. J. (2020). Age-associated changes in the response of tendon explants to stress deprivation is sex-dependent. Connective Tissue Research, 61(1), 48–62. 10.1080/03008207.2019.1648444

Cuomo, F., Kummer, F. J., Zuckerman, J. D., Lyon, T., Blair, B., & Olsen, T. (1998). The influence of acromioclavicular joint morphology on rotator cuff tears. Journal of Shoulder and Elbow Surgery, 7(6), 555–559. 10.1016/s1058-2746(98)90000-3

Dourte, L. M., Pathmanathan, L., Mienaltowski, M. J., Jawad, A. F., Birk, D. E., & Soslowsky, L. J. (2013). Mechanical, compositional, and structural properties of the mouse patellar tendon with changes in biglycan gene expression. Journal of Orthopaedic Research: Official Publication of the Orthopaedic Research Society, 31(9), 1430–1437. 10.1002/jor.22372

Durgam, S., Singh, B., Cole, S. L., Brokken, M. T., & Stewart, M. (2020). Quantitative Assessment of Tendon Hierarchical Structure by Combined Second Harmonic Generation and Immunofluorescence Microscopy. Tissue Engineering. Part C, Methods, 26(5), 253–262. 10.1089/ten.TEC.2020.0032

Farndale, R. W., Buttle, D. J., & Barrett, A. J. (1986). Improved quantitation and discrimination of sulphated glycosaminoglycans by use of dimethylmethylene blue. Biochimica Et Biophysica Acta, 883(2), 173–177.

Gögele, C., Pattappa, G., Tempfer, H., Docheva, D., & Schulze-Tanzil, G. (2025). Tendon mechanobiology in the context of tendon biofabrication. Frontiers in Bioengineering and Biotechnology, 13, 1560025. 10.3389/fbioe.2025.1560025

Gracey, E., Burssens, A., Cambre, I., Schett, G., Lories, R., McInnes, I., Asahara, H., & Elewaut, D. (2020). Tendon and ligament mechanical loading in the pathogenesis of inflammatory arthritis. Nature Reviews. Rheumatology, 16(4), 193–207. 10.1038/s41584-019-0364-x

Han, H.-M., Kim, S.-Y., & Kim, D.-H. (2025). Mechanotransduction for therapeutic approaches: Cellular aging and rejuvenation. APL Bioengineering, 9(2), 021502. 10.1063/5.0263236

Korcari, A., Przybelski, S. J., Gingery, A., & Loiselle, A. E. (2023). Impact of Aging on Tendon Homeostasis, Tendinopathy Development, and Impaired Healing. Connective Tissue Research, 64(1), 1–13. 10.1080/03008207.2022.2102004

Kwan, K. Y. C., Ng, K. W. K., Rao, Y., Zhu, C., Qi, S., Tuan, R. S., Ker, D. F. E., & Wang, D. M. (2023). Effect of Aging on Tendon Biology, Biomechanics and Implications for Treatment Approaches. International Journal of Molecular Sciences, 24(20), 15183. 10.3390/ijms242015183

Milgrom, C., Schaffler, M., Gilbert, S., & van Holsbeeck, M. (1995). Rotator-cuff changes in asymptomatic adults. The effect of age, hand dominance and gender. The Journal of Bone and Joint Surgery. British Volume, 77(2), 296–298.

Mlawer, S. J., Frank, E. H., & Connizzo, B. K. (2023). Aged tendons lack adaptive response to acute compressive injury. Journal of Orthopaedic Research, n/a(n/a). 10.1002/jor.25752

Mlawer, S. J., Pinto, F. R., Sikes, K. J., & Connizzo, B. K. (2025). Coordination of Glucose and Glutamine Metabolism in Tendon Is Lost in Aging. Journal of Orthopaedic Research: Official Publication of the Orthopaedic Research Society. 10.1002/jor.26100

Morrill, E. E., Tulepbergenov, A. N., Stender, C. J., Lamichhane, R., Brown, R. J., & Lujan, T. J. (2016). A Validated Software Application to Measure Fiber Organization in Soft Tissue. Biomechanics and Modeling in Mechanobiology, 15(6), 1467–1478. 10.1007/s10237-016-0776-3

Muljadi, P. M., & Andarawis-Puri, N. (2023). Glycosaminoglycans modulate microscale mechanics and viscoelasticity in fatigue injured tendons. Journal of Biomechanics, 152, 111584. 10.1016/j.jbiomech.2023.111584

Phillip, J. M., Aifuwa, I., Walston, J., & Wirtz, D. (2015). The Mechanobiology of Aging. Annual Review of Biomedical Engineering, 17, 113–141. 10.1146/annurev-bioeng-071114-040829

Robling, A. G., & Turner, C. H. (2009). Mechanical Signaling for Bone Modeling and Remodeling. Critical Reviews in Eukaryotic Gene Expression, 19(4), 319–338. 10.1615/critreveukargeneexpr.v19.i4.50

Rodríguez-Corrales, J.Á., & Josan, J. S. (2017). Resazurin Live Cell Assay: Setup and Fine-Tuning for Reliable Cytotoxicity Results. Methods in Molecular Biology (Clifton, N.J.), 1647, 207–219. 10.1007/978-1-4939-7201-2_14

Seitz, A. L., McClure, P. W., Finucane, S., Boardman, N. D., & Michener, L. A. (2011). Mechanisms of rotator cuff tendinopathy: Intrinsic, extrinsic, or both? Clinical Biomechanics, 26(1), 1–12. 10.1016/j.clinbiomech.2010.08.001

Sivaguru, M., Durgam, S., Ambekar, R., Luedtke, D., Fried, G., Stewart, A., & Toussaint, K. C. (2010). Quantitative analysis of collagen fiber organization in injured tendons using Fourier transform-second harmonic generation imaging. Optics Express, 18(24), 24983–24993. 10.1364/OE.18.024983

Vidal, B. de C., & Mello, M. L. S. (2019). Toluidine blue staining for cell and tissue biology applications. Acta Histochemica, 121(2), 101–112. 10.1016/j.acthis.2018.11.005

