## Supplementary figures and images for "Aged Tendons Have Impaired Mechanosensitivity and Lower Thresholds for Injury under Dynamic Compression"

### Figure S1

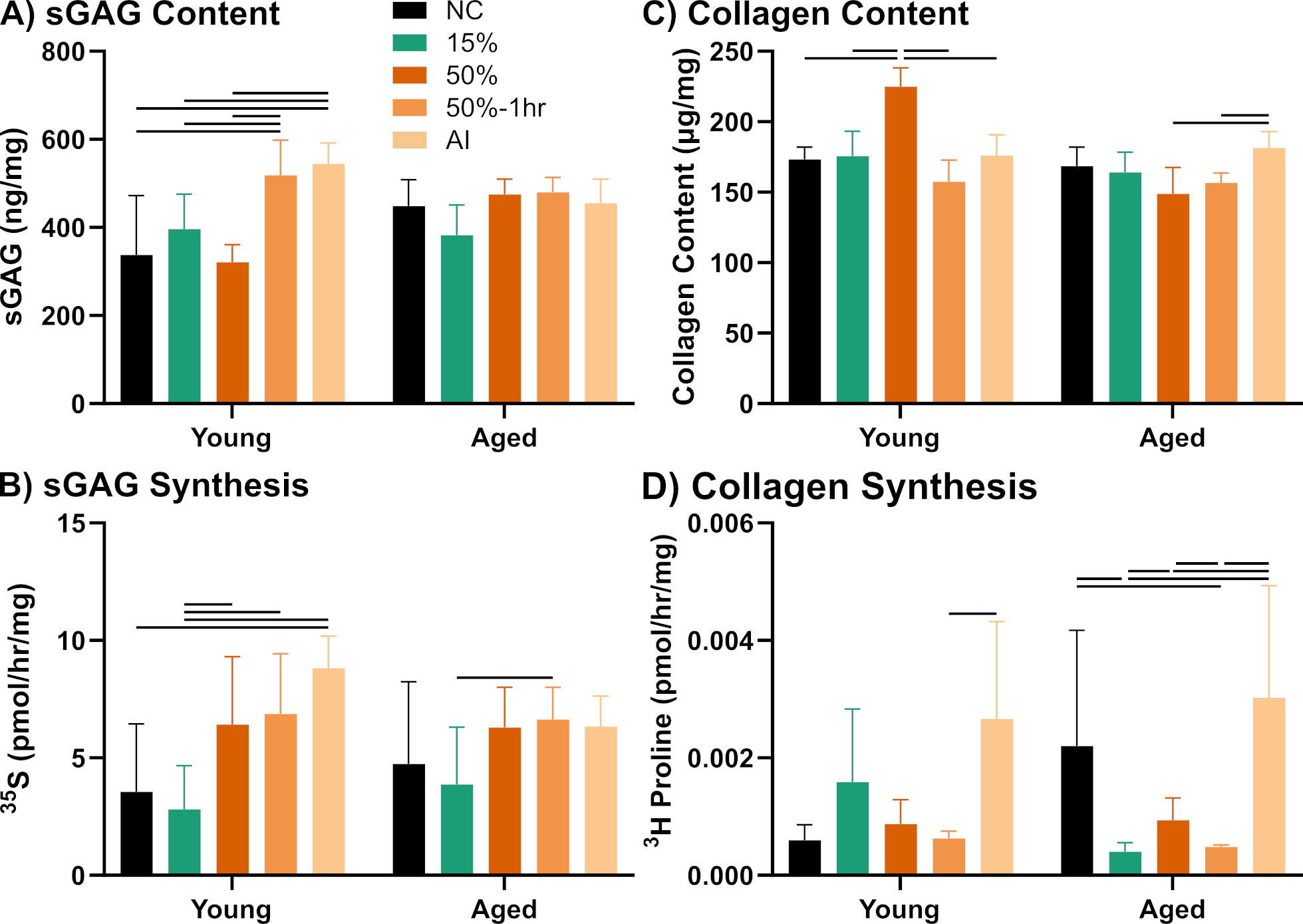
